# A multibrain advantage for cooperative human behaviour

**DOI:** 10.64898/2026.03.02.708977

**Authors:** Denise Moerel, Tijl Grootswagers, Genevieve L. Quek, Manuel Varlet

## Abstract

Collaboration is a cornerstone of human behaviour, enabling individuals to achieve complex goals that transcend individual capabilities. While previous research has largely focused on the alignment of neural representations across brains during social interactions, real-world cooperation frequently involves complementary roles, where success depends on integrating distinct yet interdependent representations. Here, we combined electroencephalography (EEG) hyperscanning and multivariate pattern analysis (MVPA) methods to investigate neural representations in pairs of participants as they jointly performed a speeded two-dimensional visual matching task with asymmetric controls to encourage task division (i.e., attending to different stimulus features). Our results provide direct neural evidence of the emergence of selective information encoding within individual brains and complementary information encoding across brains during successful cooperation. Specifically, optimally divided task dimensions give rise to a late (> 485 ms) multibrain advantage, wherein the combined neural signals of the two participants carry more target-related information than that of either brain individually. Notably, the magnitude of these information-processing modulations within and across brains robustly predicted pairs’ collective task performance. These findings have broader implications for future research on collaboration, highlighting the need to move beyond representational alignment to harness diverse human as well as machine capabilities across our societies to unlock collective potential.

## 1. INTRODUCTION

Collaboration is a ubiquitous aspect of human behaviour, enabling individuals to accomplish complex tasks that would be insurmountable alone. This is evident in various domains, including sports, military operations, and artistic performances, where individuals with diverse expertise and skills come together to achieve strategic objectives or create innovative solutions. Understanding the neural mechanisms that support such dynamic, socially coordinated behaviours in groups and teams has been advanced by hyperscanning, a technique that enables the simultaneous recording of brain activity from multiple individuals during real-time interactions (Czeszumski et al., 2020; Dumas et al., 2011; Zamm et al., 2024). Previous hyperscanning research has provided evidence of neural alignment between interacting individuals, suggesting the emergence of shared representations across brains while working toward a shared goal (Czeszumski et al., 2020; Dumas et al., 2010; Kelsen et al., 2022; Moerel, Grootswagers, Quek, et al., 2025; Mu et al., 2018; Reinero et al., 2021; Shemyakina & Nagornova, 2021). While such neural alignment can support various forms of coordinated behaviour, it does not fully capture the complexity of real-world cooperation, in which success often depends on partners with complementary roles strategically dividing task demands and encoding distinct, yet interdependent, information rather than identical information. Here, we leveraged electroencephalography (EEG) hyperscanning with multivariate decoding to investigate how information processing in pairs of participants might diverge instead of converge during visual cooperation, revealing the emergence of unique patterns of information complementarily distributed across individuals’ brains, not found in the neural activity of either individual alone and robustly predicting collective performance.

Previous research has highlighted a central role of neural and behavioural alignment in coordinated behaviours, supporting cooperation and collective performance (Coey et al., 2012; Dikker et al., 2017; Dumas et al., 2010, 2014; Garrod & Pickering, 2009; Konvalinka et al., 2011; Nalepka et al., 2019; Schilbach & Redcay, 2025; Tognoli et al., 2007). One prominent example is joint attention, the process by which individuals align their focus on the same external object, enabling shared perceptual and cognitive experiences (Bilek et al., 2015; Koike et al., 2016, 2019; Lachat et al., 2012; Saito et al., 2010). Observed already in the first years of life (Carpenter et al., 1998), engaging in joint attention has been associated with enhanced neural alignment (Koike et al., 2016; Saito et al., 2010), and has beneficial effects on cooperation and social bonding (Shteynberg, 2015). Another line of evidence comes from the joint action literature, examining how individuals coordinate their actions toward a common goal, such as moving heavy objects or playing music together (Chang et al., 2017; Keller et al., 2014; MacRitchie et al., 2017; Marsh et al., 2009; Richardson et al., 2007; Sebanz, Bekkering, et al., 2006). Neural and behavioural alignment, reflecting a range of low- to high-level sensorimotor processes (e.g., dynamical entrainment coupling, self-other mental representations) has been shown to underpin precise and flexible coordination in time and space of multiple people’s motor behaviours (Keller et al., 2014; Miyata et al., 2021; Noy et al., 2011; Richardson et al., 2007; Sebanz, Knoblich, et al., 2006; Varlet et al., 2011).

Using interbrain Representational Similarity Analysis to capture and compare the informational content in hyperscanning data, recent research has provided more direct evidence of shared representations between individuals’ brains emerging through social interaction, with new insights into the neural processes and information underpinning alignment (Moerel, Grootswagers, Quek, et al., 2025; Varlet & Grootswagers, 2024). Notably, Moerel et al. used a joint visual categorisation task to reveal interbrain information (or representational) alignment occurring in later processing stages (> 200 ms after stimulus onset). This later alignment was found to be dynamically and uniquely formed through social interaction and learning, contrasting with earlier alignment (< 200 ms after stimulus onset) remaining static over the course the experiment merely driven by shared visual input (Moerel, Grootswagers, Quek, et al., 2025). While this research provides robust evidence of shared neural representations and higher-level cognitive processes arising from social and task demands, the theoretical and methodological approach remains restricted, consistent with previous hyperscanning research (Czeszumski et al., 2020; Dikker et al., 2017; Dumas et al., 2011; Hamilton, 2021; Holroyd, 2022; Schilbach & Redcay, 2025), to the alignment of neural activity and representations. Therefore, prior research has not yet directly addressed how task-relevant information might become complementarily distributed rather than redundantly repeated across brains to harness the full potential of multi-person systems and enhance collective performance.

While direct neural evidence of information complementarily distributed between interacting brains is currently lacking, a growing evidence base from the behavioural and modelling literatures provides compelling support for its potential (Bahrami et al., 2010; Bonner et al., 2007; A. A. Brennan & Enns, 2015; Herzog & Hertwig, 2009; Hong & Page, 2004; Kameda et al., 2022; Laughlin et al., 2006; Sniezek & Henry, 1990; Stasser & Abele, 2020; Woolley et al., 2010). Studies have used a range of collective visual and decision-making tasks to show that cooperating groups can outperform even top-performing individuals, when optimally leveraging members’ diverse knowledges, capabilities, and perspectives (Bahrami et al., 2010; Bonner et al., 2007; Hong & Page, 2004; Laughlin et al., 2006; Sniezek & Henry, 1990). Importantly, effective interaction and communication within the group are critical for this collective advantage to occur, beyond an advantage simply obtained by cancelling out errors through aggregating and averaging more responses (A. A. Brennan & Enns, 2015; Herzog & Hertwig, 2009). The need for successful interaction is further evidenced by a study that revealed a stronger correlation of a groups’ collective intelligence with their members’ social sensitivity and equally shared speaking turns than with their members’ average or maximum intelligence (Woolley et al., 2010).

Crucially, to understand how such collective advantage arises at the neural level, it is necessary to investigate whether the combined brain activity of interacting individuals encodes more information than each brain does separately. To this end, it is important to extend hyperscanning research beyond alignment and shared representations, to examine the emergence of complementary representations in interacting individuals’ brains, and how they enable successful cooperation. Here we address this gap by combining hyperscanning and multivariate pattern analysis techniques to decode neural representations at both the individual and pair levels, and directly test for the emergence of selective and complementary information encoding and a “multibrain advantage” during real-time visual cooperation. We recorded EEG from twenty-five pairs of participants as they performed a speeded two-dimensional visual matching task for briefly presented target stimuli defined by the spatial frequencies of two (blue and orange) perpendicular lines (see Figure 1). On each trial, participant pairs used shared controls to select the correct option from all possible response options (25 in total presented in a 5 x 5 grid). Each participant in the pair had faster control in one cardinal direction (i.e., blue or orange dimension), enabling us to test for the selective encoding of the stimulus spatial frequency linked to the (faster) strategic control axis.

**Figure 1.**
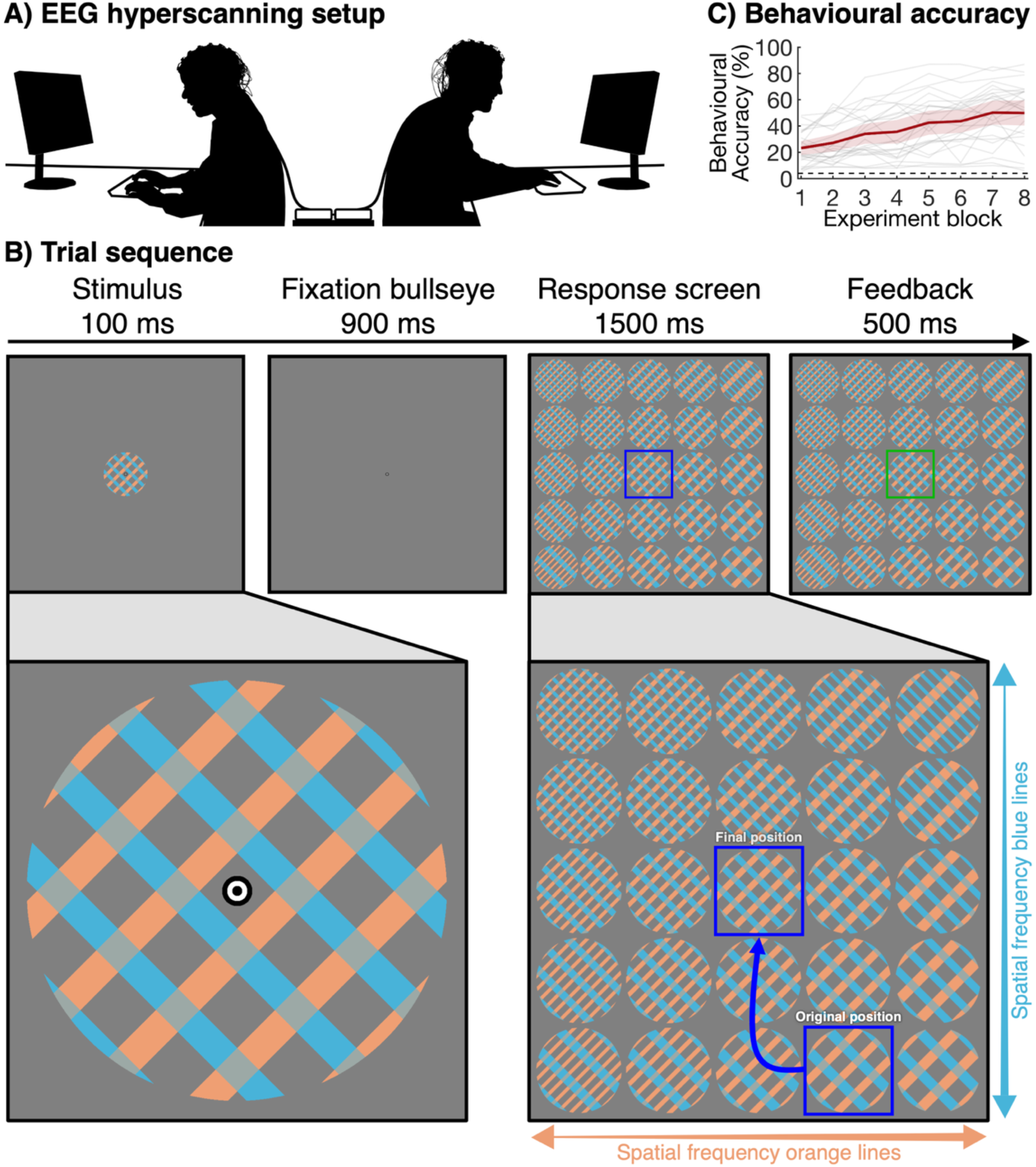
Stimuli and experiment overview. **A)** Participants in cooperating pairs were seated back-to-back in the same room, facing separate monitors that showed identical displays. **B)** Each trial contained a 100 ms target stimulus that consisted of overlaid perpendicular blue and orange lines. Lines of each colour had one of five possible spatial frequencies, such that there were 25 possible unique stimuli. The stimulus was followed by a blank screen with a fixation bullseye for 900 ms, and then by a response screen for 1500 ms that displayed all 25 response options. The participants worked together to move a blue box to select the response option that matched the previously seen target. The 25 options were arrayed such that the spatial frequency of one colour (e.g., orange) increased from left to right, while the spatial frequency of the other colour (e.g., blue) increased from top to bottom. Participant pairs received feedback after making their selection, with the blue box in the final (selected) position turning either green (correct) or red (incorrect) for 500 ms. **C)** Behavioural accuracy over the 8 experiment blocks. The grey lines show the individual pairs, and the thick red lines shows the group average. The red shaded area around the group average shows the 95% confidence interval. Chance level is 4%, as there are 25 response options, and is indicated by a black dashed line.

To anticipate our results, EEG hyperscanning decoding analyses provide direct evidence of multidimensional visual information being selectively encoded *within* individual brains, and importantly, complementarily encoded *across* individual brains during cooperation. This selective and complementary encoding increased with practice, and robustly predicted differences in collective performance between pairs. Specifically, later neural responses (> 250 ms after stimulus onset) at the level of the individual show selectively enhanced encoding of the spatial frequency associated with the strategic (faster) response dimension. In addition, we find a multibrain advantage, characterised by enhanced encoding of the stimulus target at the pair level compared to the individual level. These results provide the first direct neural evidence of complementary representations in interacting brains. In addition, they highlight the need to broaden the scope of hyperscanning research beyond alignment and synchrony to deepen our understanding of the neural processes supporting everyday human cooperation.

## 2. RESULTS

In this study, we aimed to examine whether cooperative task performance is underpinned by complementary information processing distributed across brains. Specifically, we investigated whether 1) there was evidence for task division at the behavioural and neural levels, and 2) whether there was a multibrain advantage, where the combined neural signals across brain carry more task-relevant information than individual brains alone. To this end, we recorded EEG activity from twenty-five pairs of participants using a 64 channel BioSemi ActiveTwo electrode system (BioSemi, Amsterdam, The Netherlands). The two participants in each pair were seated back-to-back in a dimly lit room, facing separate monitors (Figure 1A) while engaging in a cooperative visual task (Figure 1B). Participants viewed briefly presented stimuli composed of overlaid perpendicular blue and orange lines, with each orientation having one of five possible spatial frequencies, resulting in 25 possible unique stimuli. After a short delay, participants in each pair jointly used the arrow keys to move a box on a response grid to select the response option that matched the previously seen target stimulus. The spatial frequency for each colour varied along one axis of the response option grid (i.e., horizontal or vertical), and participants had asymmetric controls: one could move faster in the left-right dimension, the other faster in the up-down direction. After 1500 ms, the final position of the box was automatically selected. The movement constraints and time pressure were designed to encourage the participants to strategically divide the task based on colour-direction mappings. The behavioural results presented in Figure 1C show that the task was challenging, with an average accuracy around 20% (4% chance level) at the beginning of the experiment.

Performance improved over the course of the experiment with repeated practice, reaching an average accuracy of almost 50%. Notably, there was significant variability between pairs, with some showing minimal improvement.

### 2.1. Individual behavioural and neural evidence for selective information processing

Behavioural results provide strong evidence for strategic task division. We compared for each participant the percentage of responses associated with the strategic (fast) and non-strategic (slow) response dimensions as a function of experimental block (i.e., proportion of strategic responses, Figure 2B). Participants were more likely to use the strategic (fast) response buttons than would be predicted by chance (BF > 10^48^). They already preferred the strategic dimension in the first block (BF > 10^25^), but the proportion of strategic responses increased as a function of block (BF = 82.92), suggesting that participants strategically divided task demands, and their inclination and/or ability to do so increased as the experiment progressed.

**Figure 2.**
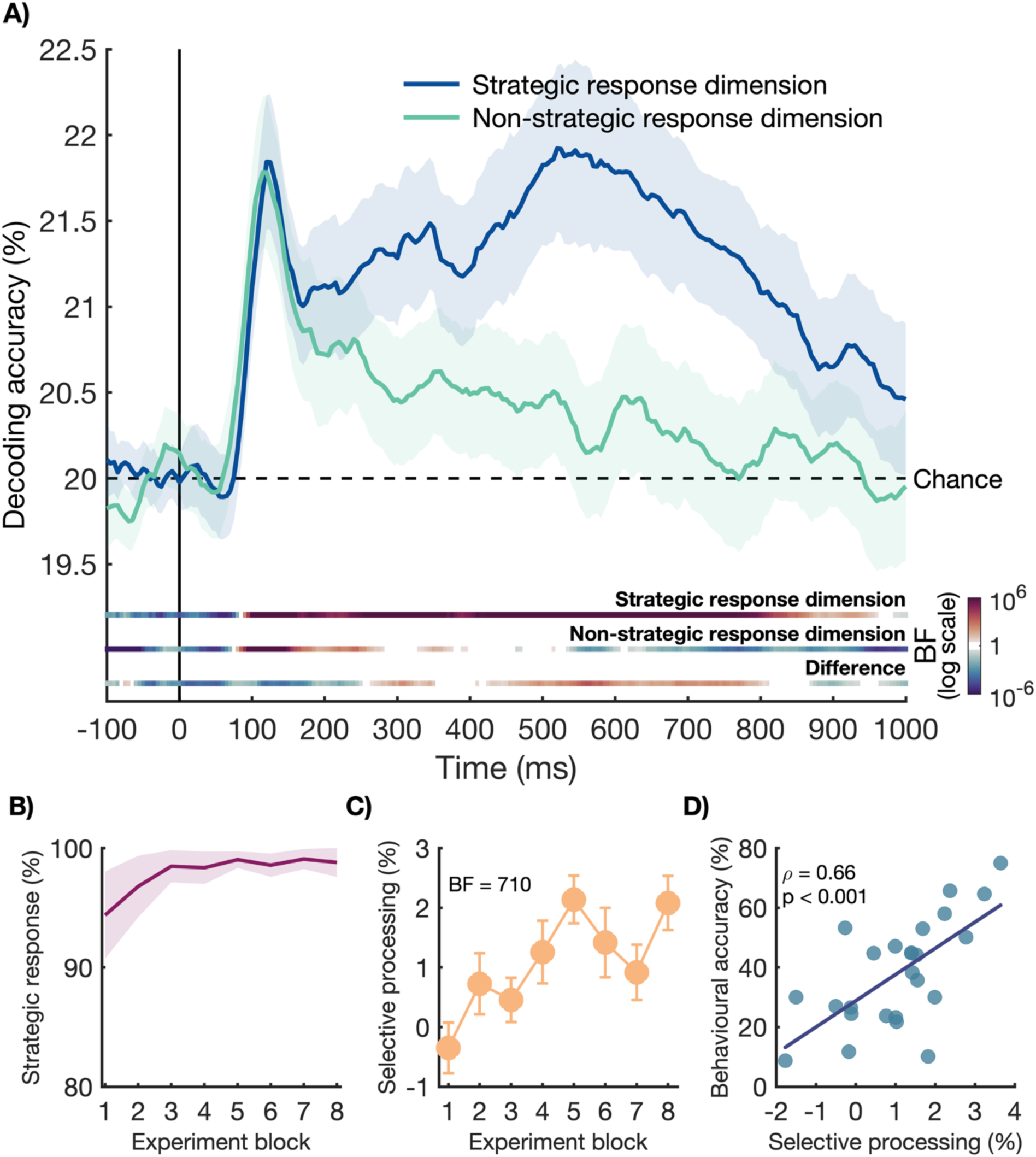
Selective information processing and its link to behavioural accuracy. **A)** The time-course of the decoding accuracy of the stimulus spatial frequency on the strategic response dimension (dark blue) and the non-strategic response dimension (green). There were 5 possible spatial frequencies on each dimension, resulting in a theoretical chance level of 20% (black dashed line). The shaded areas around the plot lines show the 95% confidence intervals. Bayes Factors are shown below the plot for the decoding accuracy associated with the strategic response dimension (top), non-strategic response dimension (middle), and the difference (strategic > non-strategic). The colours are plotted on a logarithmic scale, with red colours reflecting evidence for the alternative hypothesis, blue colours reflecting evidence for the null hypothesis, and white reflecting insufficient evidence (BF between 1/3 and 3). **B)** The percentage of responses where the participant used one of the strategic (fast) response buttons. **C)** The selective processing effect (stronger decoding of the spatial frequency on the strategic than non-strategic response dimension, averaged between 200 and 800 ms after stimulus onset) for each block in the experiment. **D)** Positive correlation between the average selective neural processing effect (averaged from 200 to 800 ms after stimulus onset across the two participants) of each pair and their average behavioural accuracy.

Importantly, this strategic behavioural division was associated with a selective enhancement of the neural encoding of the spatial frequency information linked to the strategic (fast) compared to the non-strategic (slow) response dimension (see Figure 1B). For each participant and dimension, we used a linear classifier on individual EEG data (with a 16-fold cross-validation procedure, trained and tested on 375 and 25 trials, respectively; see Methods for details) to distinguish between the 5 levels of spatial frequency at each timepoint in the epoch. Results show that individual brain activity contained information about the spatial frequencies associated with both the strategic and non-strategic response dimensions from 95 ms and 85 ms onwards, respectively. Until 245 ms, there was evidence of no difference between the decoding strength of spatial frequency information linked to the strategic and non-strategic response dimensions (BF < 0.1). However, after this time, we observed a divergence, with evidence for higher decoding accuracies for the spatial frequency information linked to the strategic response dimension from 285 ms onwards (all BF > 10, max BF = 3,852). These results indicate that overall, participants cooperated toward their shared goal (i.e., matching the stimulus target) by strategically dividing which stimulus feature to focus on, with each participant selectively attending to the dimension associated with the axis of the response screen they could more easily control.

Moreover, this selective neural enhancement of the stimulus spatial frequency of the strategic dimension was found to strengthen over the course of the experiment, in line with the behavioural results. We obtained a single measure of this selective processing effect for each block by calculating the spatial frequency decoding difference between the strategic and non-strategic response dimensions averaged from 200 to 800 ms after stimulus onset (i.e., time window with different decoding accuracy between the two dimensions; see Figure 2A). Figure 2C shows the selective processing effect across the eight experimental blocks. There was strong evidence for an increase in this neural selective processing effect over the experiment (BF = 710), mirroring the effects observed for the behavioural accuracy (Figure 1C). This suggests that the strategic collaboration between the two participants in the pair became, at both behavioural and neural levels, more efficient over time, with repeated practice.

Finally, results show that this neural selective processing effect was a robust predictor of the collective performance of the pairs (Figure 1D). We calculated a single measure of this selective processing effect for each pair by averaging the indices across the two participants, reflecting the extent to which each pair divided the task at the neural level. We also computed a single behavioural accuracy measure for each pair across the entire experiment. The results revealed a positive correlation between the pairs’ combined selective processing effect on spatial frequency decoding and behavioural accuracy (*ρ* = 0.66, p < 0.001). This suggests that pairs who strategically divided the task more strongly, by focusing their attention on the feature dimension linked to their fastest response axis, achieved higher task accuracy.

### 2.2. Multibrain advantage on the processing of target-related information

Crucially, successful cooperation was associated not only with target-related information being selectively encoded within individual brains but also complementarily encoded across brains, enabling a multibrain advantage. The combined brain activity of the two participants in the pairs contained more information about the target than each brain did separately. To test this, we computed the time-course of the target decoding within and across brains – single brain versus multibrain target decoding. Multibrain target decoding combined information carried by the neural signals from both participants in a pair, whereas single brain target decoding reflected information carried by the neural signals of each participant individually. Importantly, this analysis did not rely on combining raw channel data, but rather combined the classifier predictions, to ensure the same amount of neural data was used for both single brain and multibrain target decoding. Specifically, we took the predictions for each trial and timepoint in the epoch given by the linear classifier trained to distinguish the 5 spatial frequencies, separately for each participant and response dimension (i.e., strategic and non-strategic). We then calculated the multibrain target decoding accuracy by combining the accuracies for the spatial frequencies associated with the two participants’ strategic response dimensions within each pair (i.e., asking whether the decoding of the strategic spatial frequency for player 1 and the strategic spatial frequency for player 2 were both predicted correctly for each trial and timepoint). To obtain a comparable metric of single brain target decoding, we repeated the process, combining the accuracies for spatial frequencies associated with both the strategic and non-strategic response dimensions within each participant. This ensured that the same amount of neural decoding predictions contributed to both the multibrain and single brain decoding accuracy metrics. Finally, we also calculated a metric of non-strategic multibrain target decoding to control for a multibrain advantage that could simply arise from combining any neural decoding data from multiple individuals, potentially providing more information overall and/or reducing interference from internal and external sources, such as eye blinks or lapses in attention (Eckstein et al., 2012; Zhang et al., 2021). For this metric, we combined the accuracies for the spatial frequencies associated with the two participants’ non-strategic response dimensions within each pair.

Figure 3A shows the time-course of multibrain and single brain target decoding. There was evidence for target-related information in both the individual and combined brain signals from approximately 85 ms – 95 ms onwards. Notably, there was evidence for higher multibrain target decoding (strategic: combining the spatial frequency decoding of the two participants’ strategic response dimensions) compared to single brain target decoding from 485 ms onwards (all BF > 10, max BF = 105). This difference indicates that participants complementary encoded target-related information based on strategic task division, and that this combined encoding effort enabled a stronger representation of the target than could be achieved at the individual brain level. The comparison of strategic and non-strategic multibrain target decoding (combining strategic vs. non-strategic response dimensions) confirmed that this multibrain advantage did not trivially arise from combining multiple individuals’ neural decoding data. Multibrain target decoding was higher for combined strategic response dimensions compared to combined non-strategic response dimensions, as evidenced from approximately 464 ms onwards (all BF > 10, max BF = 291). A channel searchlight analysis showed that this later multibrain advantage – complementary target representations across brains – was largely distributed over the scalp, contrasting with early target representation, where no multibrain advantage was found, largely focused over occipital regions. These results suggest that the multibrain advantage is more than a simple early visual processing advantage but rather involves higher level visual and cognitive processes supported by more distributed neural networks.

**Figure 3.**
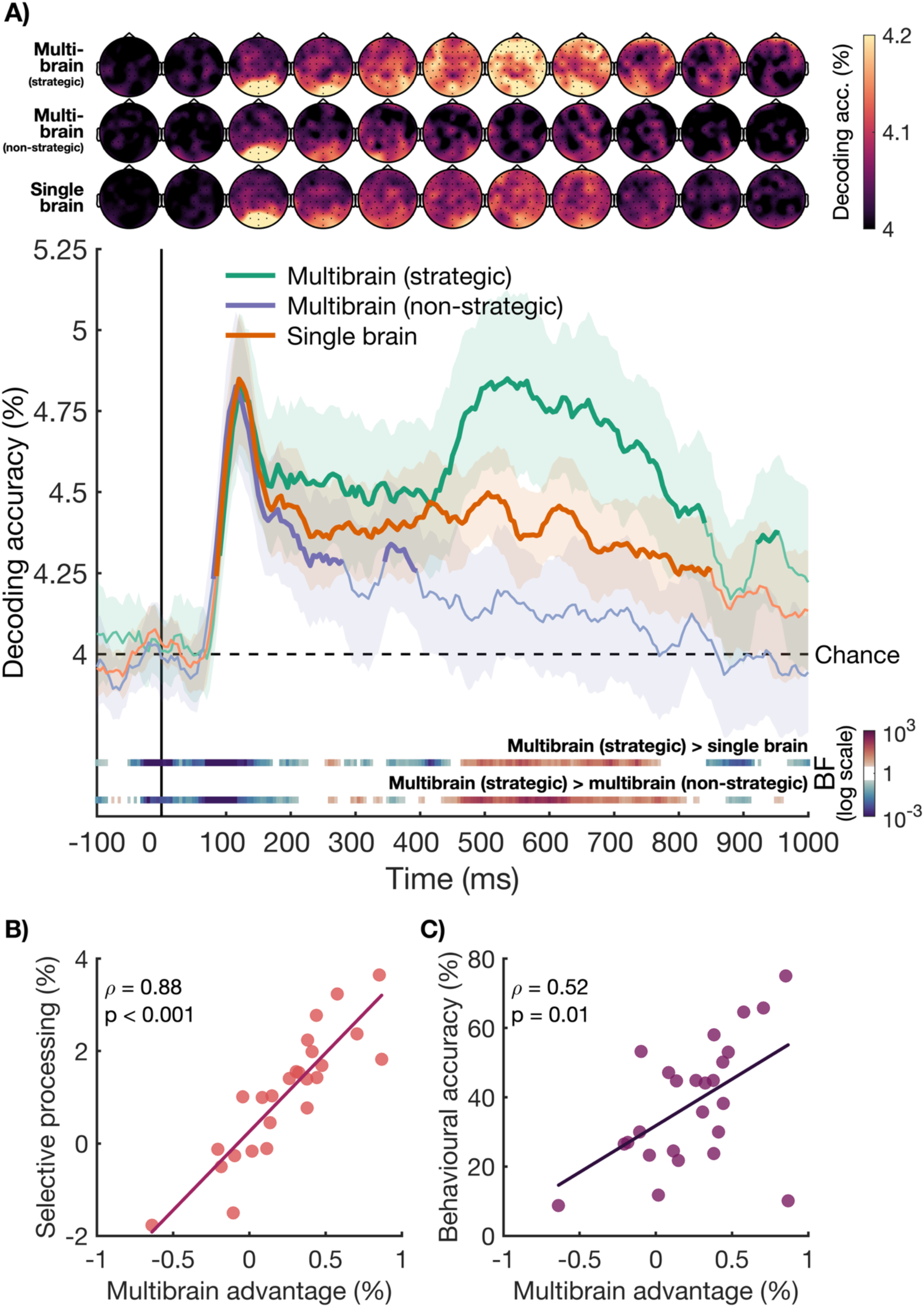
Multibrain and single brain target decoding. **A)** The time-course of target decoding accuracy based on neural data from the two participants within a pair (i.e., multibrain, green and purple) and from each participant within a pair separately (i.e., single brain, orange). The multibrain decoding accuracy represented in green combines the spatial frequency decoding of the two participants’ strategic response dimensions, and the multibrain decoding accuracy represented in purple combines the spatial frequency decoding of the two participants’ non-strategic response dimensions. Theoretical chance is at 4%, as there were 25 possible targets. The shaded areas around the plot lines show the 95% confidence interval. Thick plot lines show the timepoints where Bayes Factors are above 10, indicating strong evidence for above chance decoding. Bayes Factors for the contrasts are shown below the plot. The plot colours are based on a logarithmic scale, with red colours showing evidence for a difference, blue colours showing evidence for no difference, and white showing insufficient evidence (BF between 1/3 and 3). We used a channel searchlight to obtain topographies of decoding accuracies, shown above the decoding plot for the multibrain (top row), non-strategic multibrain (middle row), and single brain (bottom row) target decoding topographies. Lighter colours show higher decoding accuracies. **B)** The correlation between the multibrain advantage, calculated for each pair as the difference between multibrain (strategic) and single brain target decoding from 200 – 800 ms, and the neural selective processing effect, calculated for each pair as the spatial frequency decoding difference between the strategic and non-strategic response dimensions in that time window (see Figure 2). **C)** The correlation between the multibrain advantage (see Figure 3B) and the behavioural accuracy for each pair.

Finally, we show that this multibrain advantage observed across brains was strongly correlated with the selective processing effect observed within individual brains as well as the behavioural accuracy of the pairs. We found a reliable positive correlation between the multibrain advantage (strategic multibrain – single brain target decoding) and the selective processing effect (spatial frequency decoding difference between strategic and non-strategic response dimensions) averaged within the 200 – 800 ms time window (*ρ* = 0.88, p < 0.001; see Figure 3B). A strong positive correlation was also found between the multibrain advantage and behavioural accuracy (*ρ* = 0.52, p = 0.01), as depicted in Figure 3C. Together, these results demonstrate that the selective and complementary visual information encoding emerging within and across brains, respectively, during cooperation, were closely linked to each other and robustly predicted collective performance.

## 3. DISCUSSION

Our study provides new insights into the neural processes underlying cooperative information encoding, revealing how pairs of individuals strategically divide and complementarily process information to achieve shared goals. By leveraging EEG hyperscanning and multivariate pattern analysis methods, we demonstrate that cooperative task division enhances neural encoding at the pair level, giving rise to a multibrain advantage linked to better collective performance. These findings offer a new perspective on traditional views in social neuroscience, which often emphasise neural alignment and synchrony, and instead highlight the importance of divergence in neural coding for effective teamwork. With implications for optimising collaborative problem-solving, team decision-making, and human-machine interactions, our research has the potential to inform strategies for enhancing collective intelligence and performance in a wide range of real-world contexts.

### 3.1. Selective and complementary information processing during cooperation

Using EEG hyperscanning during real-time cooperation, we provide strong neural evidence of task division, with selective information encoding within brains and complementary information encoding between brains, highlighted in the form of a multibrain advantage. The processing of the two dimensions of the visual task used here (spatial frequencies of the blue and orange lines) was strategically divided by participants to facilitate collective performance. We observed robust positive correlations between the selective and complementary information encoding at the neural level and the pairs’ behavioural accuracy. The stronger the selective and complementary information encoding (i.e., selective processing index within individual brain activity and multibrain advantage combining the two participants’ brain activity), the better the pairs were at finding and matching the stimulus target. Such cooperative information encoding dynamically emerges from the interaction between the task constraints, including time-pressured visual processing and response, and the participants’ movement constraints on the two dimensions. Cooperative information encoding was not instantaneous but emerged and strengthened, together with behavioural accuracy, through practice.

Our findings underscore the non-trivial nature of cooperative information encoding, requiring effective interaction and communication within groups for resources to be optimally coordinated across people: Despite the relatively straightforward two-dimensional design of our task, not all pairs were able to successfully implement the optimal solution. This was not only reflected in poor behavioural accuracy over the course of the experiment but also in weaker cooperative information encoding (i.e., selective processing within brains and multibrain advantage) than other pairs. Moreover, our results show it is not just about bringing more people and brains together – not all multibrain combinations led to an advantage – but rather optimally leveraging individual capabilities, constrained in our experiment by movement speed on the different axes. These findings provide a novel neuroscientific perspective on previous behavioural and modelling research, including that focused on the ability of groups to outperform individuals (Bahrami et al., 2010; Bonner et al., 2007; Hong & Page, 2004; Laughlin et al., 2006; Sniezek & Henry, 1990). While our well-controlled two-dimensional visual task cannot fully capture the diversity and complexity of everyday cooperation, our results already emphasise the inherent variability of collective strategies and multibrain advantages that can emerge in groups. Previous research has shown that a range of efficient behavioural divisions are adopted when the experimental paradigm allows for multiple ways to divide the task (Andrade-Lotero & Goldstone, 2021; S. E. Brennan et al., 2008). Multibrain methods can play a key role in uncovering the neural underpinnings of such unique collective coordination strategies in future research.

### 3.2. Underlying neural processes of cooperative information processing

The time-courses of the selective processing and multibrain advantage effects, depicted in Figures 2 and 3, suggest dynamic modulations in participants’ neural activity associated with later visual processing stages. Early neural responses before 200 ms appear largely stable, while later neural responses seem to reflect more directly participants’ adaptation to task and social demands. This is consistent with the involvement of higher-level visual and cognitive processes supporting selective and complementary target representations. These findings align with recent research that examined joint visual categorisation with hyperscanning and revealed information alignment emerging between participants’ neural representations at later processing stages, suggesting the involvement of processes such as attention, visual categorisation, and decision-making (Moerel, Grootswagers, Quek, et al., 2025). More generally, these results are in line with a larger body of evidence from single-participant studies that showed feature-based attention effects on the processing of simple visual stimuli from 200 ms after stimulus onset onwards (Moerel et al., 2022, 2024; Smout et al., 2019).

The involvement of such cognitive processes is consistent with the topographical distributions of the multibrain advantage shown in Figure 3, which indicate early target decoding distributed over occipital regions, in contrast to later target decoding advantage, which is more distributed across the scalp. This might reflect activity from frontoparietal regions, that have been associated with higher-level information processing and decision-related mechanisms supporting behavioural selection and control (Jackson et al., 2016; Jackson & Woolgar, 2018; Löffler et al., 2019; Moerel et al., 2024). Neuroimaging techniques with higher spatial resolution, such as functional magnetic resonance imaging (fMRI), might enable future research to gain further insight into the brain regions underpinning these later neural modulations and cooperative information encoding. Brain stimulation techniques might also help test more directly the causal role of these regions (Novembre & Iannetti, 2021; Varlet et al., 2017). Together, these findings build growing evidence of later flexible neural responses dynamically responding to task and social demands, enabling both selective and complementary information processing during cooperation.

### 3.3. New perspectives for hyperscanning research

Showing a central role of complementary information processing, our findings underscore the critical need to expand hyperscanning research beyond alignment to capture the full range of the neural processes supporting everyday social interactions. Our research extends prior studies on joint attention, which have largely focused on aligned attention between individuals, by emphasising the equally important role of complementary attention (Bilek et al., 2015; Koike et al., 2016, 2019; Lachat et al., 2012; Saito et al., 2010). It also builds upon joint action literature that examines shared representations and associated behavioural and neural alignment enabling individuals coordinating their actions with each other (Dumas et al., 2010; Gugnowska et al., 2022; Keller et al., 2014; Miyata et al., 2021; Sebanz, Bekkering, et al., 2006; Sebanz, Knoblich, et al., 2006; Varlet et al., 2020). Our results demonstrate that in cooperative settings with complementary roles, neural representations of the same visual input can diverge rather than align, and that this strategic information division at the neural level can enhance group performance towards shared goals. Successful cooperation does not necessarily require individuals to represent the same information; instead, it can benefit from distributing sensorimotor and cognitive processes across individuals. This represents a significant shift from the dominant focus in hyperscanning research, and social neuroscience more generally, on alignment and synchrony as the primary signatures of collaboration (Djalovski et al., 2021; Garrod & Pickering, 2009; Hamilton, 2021; Holroyd, 2022; Novembre & Iannetti, 2021; Reinero et al., 2021; Zamm et al., 2024). Our results emphasise the value of divergence in neural coding, suggesting that coordinated differences can drive effective teamwork, which needs to be integrated into hyperscanning theories and methods to fully capture the neural processes supporting social interaction beyond alignment and synchrony. Importantly, our findings also highlight the potential of multivariate pattern analysis methods for hyperscanning research and moving beyond alignment. Notably, most hyperscanning research has focused on interbrain synchrony measures (Czeszumski et al., 2020; Dikker et al., 2017; Dumas et al., 2010, 2011; Pick et al., 2024; Reinero et al., 2021; Zamm et al., 2024) with the aim of capturing neural alignment, which have limitations because they do not allow direct indexing of the information contained in neural signals. Multivariate pattern analysis methods increasingly appear as a powerful alternative to capture and compare more directly neural representations between individuals and reduce confounds from shared sensory input (Anders et al., 2011; Konvalinka & Roepstorff, 2012; Moerel, Grootswagers, Chin, et al., 2025; Moerel, Grootswagers, Quek, et al., 2025; Varlet & Grootswagers, 2024; Zada et al., 2024). Here, we further underscore this potential with high temporal resolution tracking of the information contained in neural signals at both individual and collective levels. Crucially, this new multibrain approach enabled us to move beyond the neural encoding of redundant information across individuals and instead focus on complementary neural encoding through strategic task division. It revealed information present in the combined neural signals of the pair, surpassing information contained in individual neural signals. The potential benefit of multibrain information decoding has been suggested in previous studies that combined brain activities recorded separately or in non-interactive settings (Eckstein et al., 2012; Zhang et al., 2021). Our study demonstrates a multibrain information decoding advantage during real-time cooperation emerging from effective interaction and communication within groups, reflecting optimally distributed cognitive resources across people and robustly predicting their collective behavioural performance. This new multibrain approach, leveraging hyperscanning and multivariate pattern analysis methods, has critical importance for future research, not only to move beyond alignment, but also to understand at the informational level the neural processes underpinning collective intelligence and performance.

To conclude, using multivariate decoding on EEG hyperscanning data, we provide the first direct neural evidence of information processing complementarily distributed across brains during real-time visual cooperation. We showed that pairs of participants who optimally cooperated encoded more target-related information collectively than either individual alone and this advantage robustly predicted their collective performance. These findings demonstrate that cooperative task division enhances neural encoding at the pair level, yielding a measurable multibrain advantage. This novel multibrain approach provides a powerful framework for investigating real-time social interactions, moving beyond interbrain alignment and synchrony to directly assess neural information at the team level. Its relevance extends to today’s culturally diverse and interconnected societies, where successful collaboration hinges on integrating varied viewpoints and expertise. Additionally, it has critical importance for human-machine systems becoming ubiquitous in our societies, where capabilities must be complementarily coordinated to enhance performance. Ultimately, our findings highlight the significance of considering the multibrain basis of human collaboration, which will be vital for optimising collective performance as we continue to develop more complex social and technological systems.

## 4. METHODS

### 4.1. Participants

Fifty-two participants, grouped into twenty-six pairs, took part in this experiment. One pair did not finish the experiment, and was excluded from the analysis, resulting in a final sample of twenty-five pairs (mean age = 23.92 years, SD = 3.74 years, range = 18 – 38 years, 30 female/20 male, 44 left-handed/6 right-handed). All participants reported normal or corrected-to normal vision. All but one participant reported normal colour vision, with one participant reporting red-green colour blindness. The study was approved by the Western Sydney University ethics committee (H14498), and participants provided written and verbal informed consent before participating. The EEG session, including setup, took approximately 120 minutes to complete, and participants received 60$ AUD for their participation.

### 4.2. Stimuli and EEG experiment

During the experiment, the two participants in the pair were seated back-to-back, facing separate monitors, in a dimly lit room (Figure 1A). We collected EEG data from both participants while they engaged in the following task (Figure 1B). Participants saw blue and orange lines, presented in a circular window, overlaid at fixation for 100 ms, followed by a blank screen with a central bullseye for 900 ms. After a delay, the participants saw a response screen for 1500 ms, showing 25 possibilities and a movable box. The two participants in the pair had to work together to move the box to the correct response option, that reflected the stimulus they saw.

All stimuli were presented on a mid-grey background with a refresh rate of 120 Hz via a VIEWPixx/EEG monitor (VPixx Technologies Inc., Saint-Bruno, Ontario, Canada), using the Psychopy library (Peirce et al., 2019) in Python. Participants were seated approximately 60 cm from the screen. The stimuli measured 200 by 200 pixels (approximately 5.06 degrees of visual angle) and were overlaid with a central bullseye measuring 18 by 18 pixels (approximately 0.46 degrees of visual angle). Participants were instructed to keep fixation on the bullseye. The stimuli were based on those used in previous work to investigate selective processing effects in single participants (Moerel et al., 2022, 2024), and consisted of blue (RGB = 239, 159, 115) and orange (RGB = 72, 179, 217) lines, that were approximately equiluminant. The blue lines were always tilted 45° counter-clockwise of vertical, and the orange lines were always tilted 45° to clockwise of vertical, resulting in an offset of 90° between the lines. There were 25 unique stimuli, obtained by combining 5 possible spatial frequencies, 3, 4, 5, 7, or 9 cycles per stimulus, separately for the orange and blue lines.

The response screen showed all 25 unique stimuli and was always ordered in such a way that moving right to left changes the spatial frequency of one colour, whereas moving up and down changes the spatial frequency of the other colour (for an example, see Figure 1B). The spatial frequencies were either ordered from highest to lowest, or lowest to highest, separately for blue and orange, making four possible response screens in a block. Which colour was linked to left-right, and which colour to up-down changed each block, but remained the same within a block. The four possible response screens in a block were randomly presented, which means there could be no meaningful motor preparation until this screen was presented. The response screen therefore ensured that there was minimal contamination of motor preparation or execution in the EEG signal. During this screen, participants had 1500 ms to move to box to the correct position, matching the stimulus they saw. Participants saw the same screen, which means they could both move the same box and observe the movements from their partner. They both used four keys on the keyboard, labelled with arrows, to make their response. Participants could move the box in all four directions, but for each participant, two of the four buttons moved the box instantly (i.e., on the next screen refresh), whereas the other two buttons moved the box more slowly, taking approximately 200 ms to get to the neighbouring stimulus. Critically, for one of the two participants the up and down buttons worked fast, while for the other participant the left and right buttons worked fast. This stayed the same throughout the experiment. The response time-out of 1500 ms added time pressure, encouraging participants to use strategies to be successful at this task. The consistency in which colour was linked to which movement dimension ensured that it was possible to divide the task based on the colour/orientation of the stimuli. That is, the participant with the fast left-right buttons could focus on the colour that was linked to the left-right dimension, and the participant with the fast up-down buttons could focus on the colour that was linked to the up-down dimension. The start position of the box was random, with the only constraint that it could not be the same as the correct position. Once the time-out of 1500 ms was reached, the current position of the box was automatically locked in as the response. This was followed by feedback for 500 ms. During the feedback, the response screen with the box in its final selected position was presented, with the colour of the box presented in either green or red, depending on whether the response was correct or incorrect, respectively. This was followed by a blank screen with a central bullseye for 1000 ms between trials.

Participants completed 8 blocks, with 100 trials per block. Each block consisted of 4 repeats of the 25 unique stimuli. The stimuli were presented in a random order. Which colour was linked to which movement dimension (i.e., horizontal or vertical) changed for each block. In between blocks, participants received feedback of their score for that block (% of correct trials) and were given the opportunity to have a break.

### 4.3. EEG acquisition and pre-processing

We collected continuous EEG data from 64 channels, digitised at a sampling rate of 2048 Hz, using the BioSemi Active-Two electrode system (BioSemi, Amsterdam, The Netherlands). Electrode placement followed the international 10-20 system (Oostenveld & Praamstra, 2001). Data pre-processing was conducted in MATLAB using the FieldTrip toolbox (Oostenveld et al., 2011). The data were re-referenced to the average of all channels, with filters applied: a high-pass filter at 0.1 Hz, a low-pass filter at 100 Hz, and a 50 Hz band-stop filter to remove line noise. We down-sampled the data to 200 Hz and segmented it into epochs ranging from −100 ms to 1000 ms relative to stimulus onset. The selected epoch excludes the part of the trial where the response screen was presented. No meaningful motor response could be prepared during the selected epoch as the response screen was presented in a pseudo-random order, thus preventing any motor related effects. A baseline correction was performed using the interval between −100 and 0 ms.

### 4.4. Behavioural analyses

We calculated a combined behavioural accuracy for each pair, separately for each block, by determining the percentage of trials where the correct target was selected. As the target was defined as the correct combination of spatial frequency for both line colours, the chance level was 4%. To quantify whether there was an improvement in task performance over time, we fit a linear function to the accuracy over blocks for each pair and obtained the slopes. We then determined whether there was evidence that the slopes were above 0.

In addition, we determined which response buttons participants used, to gain insight into whether they were dividing the task. For each individual participant and block, we determined the percentage of responses that were associated with the fast response buttons.

### 4.5. EEG decoding analyses

#### 4.5.1. Single brain spatial frequency decoding analysis

We used multivariate decoding on time-varying EEG data (Grootswagers et al., 2017) to determine if and when there was information about the spatial frequency associated with 1) the strategic response dimension (fast response buttons) and 2) the non-strategic response dimension (slow response buttons). We ran the same analysis separately for the orange and blue lines and then collapsed the decoding accuracies across line colours. We used a regularised Linear Discriminant Analysis classifier, with a 16-fold cross-validation, implemented using the CoSMoMVPA toolbox (Oosterhof et al., 2016). This classifier was chosen as it provides a good balance between computational requirements and performance (Grootswagers et al., 2017; Guggenmos et al., 2018). The decoding analysis was performed on the raw activation values across the 64 EEG channels. For each fold, we trained the classifier on 375 trials to distinguish the 5 different spatial frequencies and tested on the remaining 25 trials. Importantly, the blue and orange lines were orthogonal dimensions in the design, following previous work (Moerel et al., 2022, 2024), which means we were able to assess the feature decoding of the lines of one colour without contributions from the lines of the other colour.

We further assessed whether the selective processing effect, calculated as the difference in spatial frequency decoding accuracy associated with the strategic compared to non-strategic response dimension, emerged over time in the experiment. We used the same decoding analysis as described above but calculated the time-course of spatial frequency decoding accuracy separately for the features associated with the strategic and non-strategic dimension for each of the 8 blocks in the experiment. We then obtained a single measure of the selective processing effect for each block by calculating the difference in spatial frequency decoding between the features associated with the different response dimensions, averaged across the time-window between 200 and 800 ms relative to stimulus onset. This window was chosen to select the timepoints that yielded evidence for statistical differences in decoding accuracy between the two features (all BF > 10), which were associated with the strategic compared to non-strategic response dimension.

Finally, we determined whether there was a link between the neural selective processing effect on spatial frequency decoding and behaviour. For each pair of participants, we calculated a single measure of the neural selective processing effect on spatial frequency decoding by calculating the difference in spatial frequency decoding associated with the strategic compared to non-strategic response dimension, averaged across the time-window between 200 and 800 ms relative to stimulus onset and across all blocks. We did this separately for the two individuals in the pair and then averaged the neural selective processing effect measures across individuals to obtain a single measure per pair. The behavioural accuracy measure per pair was calculated as the percentage of correct responses across the experiment. We then calculated a Pearson correlation between the neural selective processing effect on spatial frequency decoding and behaviour.

#### 4.5.2. Multibrain target decoding analysis

We used multibrain decoding (Eckstein et al., 2012; Zhang et al., 2021) to test whether the combined neural signals from the two participants, measured through EEG, captured more information about the stimulus target than those from either participant alone. We used the same classification analysis as described above, decoding each stimulus feature at the individual participant level, and then combining them into a measure of multibrain target decoding. First, we obtained at the individual participant level the predicted spatial frequency associated with the strategic colour for each timepoint in the epoch. We did this separately for the two participants in the pair and then combined the two into multibrain decoding by calculating whether the feature associated with the strategic response dimension for both individuals in the pair was correctly classified. By combining the decoding of the spatial frequency for both the orange and blue lines, we obtain the target decoding, where the chance level is 4%. We did this separately for each timepoint and each trial and then calculated the mean decoding accuracy across all trials.

We compared the multibrain target decoding to single brain target decoding to determine whether there was a multibrain advantage (Eckstein et al., 2012). For this, we used the predicted spatial frequency associated with both the strategic and non-strategic colour for each individual trial, timepoint, and participant. We calculated the target decoding accuracy by determining whether the spatial frequencies associated with both the strategic and non-strategic dimensions were classified correctly, averaged across individual trials. To determine whether an advantage of multibrain compared to single brain decoding could be driven simply by using the signal of two brains compared to one, for example by increasing the amount of available data (Eckstein et al., 2012; Zhang et al., 2021), or reducing internal and external interference, such as eye blinks or attention lapses (Zhang et al., 2021), we also calculated a multibrain decoding accuracy based on the non-strategic response dimensions of the two participants of the pair. Finally, to determine the topographical distribution of target decoding at the single brain and multibrain level (based on the strategic and non-strategic response dimensions), we ran a searchlight across the 64 channels. For each channel, we selected 4-5 neighbouring channels and repeated the same analyses described above. This resulted in a topography of target decoding accuracies.

To test whether an observed multibrain advantage could be linked to participants dividing the task and each attending to different spatial frequencies, we correlated the multibrain advantage with the neural selective processing effect for each pair. We used the same measure of the neural selective processing effect on spatial frequency decoding as described above, by calculating the difference in spatial frequency decoding associated with the strategic compared to non-strategic response dimension, averaged across the time-window between 200 and 800 ms after stimulus onset. We calculated for each pair the multibrain advantage as the difference between the multibrain target decoding and the average single brain target decoding in the same time window. We then calculated a Pearson correlation between the multibrain advantage in target decoding and neural selective (spatial frequency decoding) processing effect. Finally, we calculated a Pearson correlation between the multibrain advantage and the behavioural accuracy to test the link between the two.

### 4.6. Statistical inference

We used Bayesian statistics to evaluate the evidence for the null hypothesis (chance decoding) and the alternative hypothesis (above-chance decoding) at each timepoint (Morey & Rouder, 2011; Rouder et al., 2009; Teichmann et al., 2022), with the Bayes factor package in R (Morey & Rouder, 2018). We performed a Bayesian *t*-test at each timepoint, using a half-Cauchy (directional) prior centred at (*δ* = 0) with the default prior width of 0.707. We excluded effect sizes between *δ* = 0 and *δ* = 0.5 (Teichmann et al., 2022). We also assessed whether the decoding accuracy differed between 1) the spatial frequency associated with the strategic compared to the non-strategic response dimension, 2) the multibrain (strategic, based on the strategic dimensions) compared to single brain target decoding, and 3) the multibrain based on the strategic compared to non-strategic dimensions. We obtained the difference between the decoding accuracies and tested against 0, using the same Bayesian *t*-test as described above for each timepoint. Because we had hypotheses about the direction of the effect (strategic spatial frequency > non-strategic spatial frequency, multibrain (strategic) > single brain, and multibrain (strategic) > non-strategic multibrain), we used half-Cauchy priors to make the test directional. Bayes factors (BFs) below 1 indicated evidence for the null hypothesis (chance decoding or no difference), while values above 1 indicated evidence for the alternative hypothesis (chance decoding or a difference). We defined the onset of above chance decoding or difference by finding the first three consecutive timepoints where all Bayes Factors were >10.

Finally, we used a Bayesian t-test to determine whether there was evidence that the slopes, fitted to the behavioural task performance for each pair over time in the experiment, were above 0. We used the same method described above, implementing a half-Cauchy prior to test for an increase in task accuracy only, without a null interval.

## ACKNOWLEDGMENTS

This work was supported by an Australian Research Council (ARC) Discovery Project awarded to M.V. (DP220103047) and an ARC Discovery Early Career Researcher Award awarded to T.G. (DE230100380).

## 5. DATA AND CODE AVAILABILITY

The dataset will be made available upon publication.

